# OSCAR: Optimal subset cardinality regression using the L0-pseudonorm with applications to prognostic modelling of prostate cancer

**DOI:** 10.1101/2022.06.29.498064

**Authors:** Anni S. Halkola, Kaisa Joki, Tuomas Mirtti, Marko M. Mäkelä, Tero Aittokallio, Teemu D. Laajala

## Abstract

In many real-world applications, such as those based on patient electronic health records, prognostic prediction of patient survival is based on heterogeneous sets of clinical laboratory measurements. To address the trade-off between the predictive accuracy of a prognostic model and the costs related to its clinical implementation, we propose an optimized *L*_0_-pseudonorm approach to learn sparse solutions in multivariable regression. The model sparsity is maintained by restricting the number of nonzero coefficients in the model with a cardinality constraint, which makes the optimization problem NP-hard. In addition, we generalize the cardinality constraint for grouped feature selection, hence making it possible to identify key sets of predictors that may be measured together in a kit in clinical practice. We demonstrate the operation of our cardinality constraint-based feature subset selection method, named OSCAR, in the context of prognostic modelling of prostate cancer, where it enabled one to determine the key explanatory predictors at different levels of model sparsity, and to explore how the model sparsity affects the model accuracy and implementation cost.

**Author summary:** Feature selection has become a crucial part in building biomedical models, due to the abundance of available predictors in many applications, yet there remains an uncertainty of their importance and generalization ability. Regularized regression methods have become popular approaches to tackle this challenge by balancing the model goodness-of-fit against the increasing complexity of the model in terms of coefficients that deviate from zero. Regularization norms are pivotal in formulating the model complexity, and currently *L*_1_ (LASSO), *L*_2_ (Ridge Regression) and their hybrid (Elastic Net) norms dominate the field. In this paper, we present a novel methodology using the *L*_0_-pseudonorm, also known as the best subset selection, which has largely gone overlooked due to its challenging discrete nature. Our methodology makes use of a continuous transformation of the discrete optimization problem, and provides effective solvers implemented in a user friendly R software package. We exemplify the use of *oscar-package* in the context of prostate cancer prognostic prediction using both real-world hospital registry and clinical cohort data. By benchmarking the methodology against related regularization methods, we illustrate the advantages of the *L*_0_-pseudonorm for better clinical applicability and selection of grouped features.

## Introduction

Current cancer incidence is more than 19 million new cases per year and rapidly rising globally [1]. Despite the successful development of medical treatments that have decreased the mortality of cancer patients, cancer remains one of the most common causes of death, thus leading to dire need for more precise and prognostic insights into patient care. Prognostic prediction is fundamental in patient management, since it enables the assessment of prognosis in diagnostic phase and prediction of the course of the disease for an individual patient after treatment or disease relapse. Predicting the risk of cancer recurrence or death, based on the individual patient characteristics and laboratory measurements, helps to understand, which patients would benefit from a standard treatment and which are better assigned to palliative care or treated with alternative therapy regimens. In clinical practice, survival prediction is typically done based on laboratory tests, which are many times numerous and thus expensive. From an economical point of view, prognostic modelling should be both accurate and cost-effective, and the prognostic models should not become too complex to enable clinical implementation. In this particular aspect, feature selection strategies, such as regularization in regression modelling, play a key role.

Prostate cancer is one of the most common cancers diagnosed in men and among the top causes of cancer mortality [1]. Although the prognosis of prostate cancer is generally good, a considerable number of patients either have a metastasized disease at the time of diagnosis or they develop a potentially lethal recurrent disease during follow-up after the initial treatment. Prostate-specific antigen (PSA) is currently considered as the default marker of disease progression during the follow-up. However, when prostate cancer develops into a hormonal treatment independent state (i.e. castration resistant prostate cancer), more rigorous testing including additional markers is needed for more accurate patient stratification [2]. Given the high prevalence of prostate cancer globally, it is not trivial to consider the costs of this testing during follow-up, further increasing the need for cost-effective modelling strategies.

Risk classification models for prostate cancer are traditionally applied either in diagnostic phase or primary treatment phase. Most current prognostic models contain Gleason score, which is considered the most significant factor for early disease course estimation [3]. In contrast, our objective here was to make prognostic prediction of patients who have already developed metastatic castration-resistant prostate cancer, and therefore seek to investigate prognostic features beyond Gleason score. Regularized Cox regression models have been a popular choice for such prognostic modelling purposes [4–8]. For example, in the DREAM 9.5 Prostate Cancer Prediction Challenge [6], our top-performing model was based on an ensemble of regularized models with Cox regression [9].

In the present work, the prognostic modelling framework for prostate cancer is also based on the Cox’s proportional hazards model [5,10]. The base Cox’s model is extended by introducing a novel feature selection regularization strategy. To this end, we use a cardinality constraint expressed with the *L*_0_-pseudonorm to restrict the number of nonzero coefficients. Including the cardinality constraint complicates the optimization, since this constraint is discontinuous and nonconvex, which makes the problem NP-hard (nondeterministic polynomial hard) [11]. Due to the NP-hard optimization problem, there has been a lack of implementations utilizing this modelling strategy. Some modelling approaches with *L*_0_-implementations, such as [12,13], have been developed for generalized linear models, such as linear and logistic regression, but they do not offer solutions for the Cox model essential for prognostic predictions. To the best of our knowledge, there is only one *L*_0_-implementing Cox’s proportional hazards model, the augmented penalized minimization-*L*_0_ (APM-*L*_0_) [14], which approximates the *L*_0_ approach, and iterates between a coordinate descent based convex regularized regression and a simple hard-thresholding estimation.

Our implementation differs from the approach of APM-*L*_0_. First, we rewrite the cardinality constraint with its exact DC (Difference of two Convex functions) representation after which the constraint is added to the objective function utilizing a penalty function approach [15]. This leads to a continuous nonsmooth objective function. However, the nonconvexity remains even after the transformation. In our method, the optimization is done with two sophisticated solvers: the double bundle method (DBDC) [16,17] for DC optimization and the limited memory bundle method (LMBM) [18,19] for nonsmooth large-scale optimization. Both solvers are capable of handling the exact DC representation of the cardinality-constrained problem after it has been transformed into a penalty function form. In addition to the advanced optimization methods and inclusion of the cardinality constraint, we generalize the cardinality constraint to also control the number of used kits linking predictors that come with the same cost together. Instead of a single measurement, in practice, many features are often measured together as kits (such as complete blood count). In our method, such kit structure can be included, thus enabling the selection of relevant predictor sets instead of just single predictors.

In this work, we present a new *L*_0_ regularization method OSCAR (Optimal Subset CArdinality Regression) and exemplify it with Cox’s proportional hazards model in prognostic prediction of prostate cancer. In addition to survival prediction, the OSCAR method implements the binomial model for logistic regression problems and the linear regression model with mean square error (see e.g. [20]). The OSCAR method is tested in four separate training data cohorts: TYKS (real-world hospital registry data) [8], and VENICE, MAINSAIL and ASCENT (randomized clinical trials) [6]. We use bootstrap (BS) and cross-validation (CV) analyses to ensure generalization ability of the model. The model performance accuracy is investigated alongside the corresponding predictor costs; this helps to identify which models are cost-effective (i.e., max accuracy, min cost). Combining these two objectives makes the underlying problem a multi-objective optimization problem. We note that the process of fitting the Cox’s proportional hazards model (i.e. accuracy) for all the required cardinalities is one way to obtain an approximation of the Pareto-front [21] in this multi-objective problem. These Pareto-fronts can then be provided for the end-users for domain-expert driven decision making. Finally, the models selected based on the Pareto-fronts are also tested in the validation cohorts independent from the training data sets separated before model fitting. In OSCAR, we refer to cardinality as the number of predictors or groups of predictors (i.e. kits) in the model. Schematic illustration of the OSCAR method is presented in Fig 1. In addition to OSCAR analyses, we compare the results to traditional LASSO [4] and *L*_0_-augmented APM-*L*_0_ [14].

**Fig 1.**
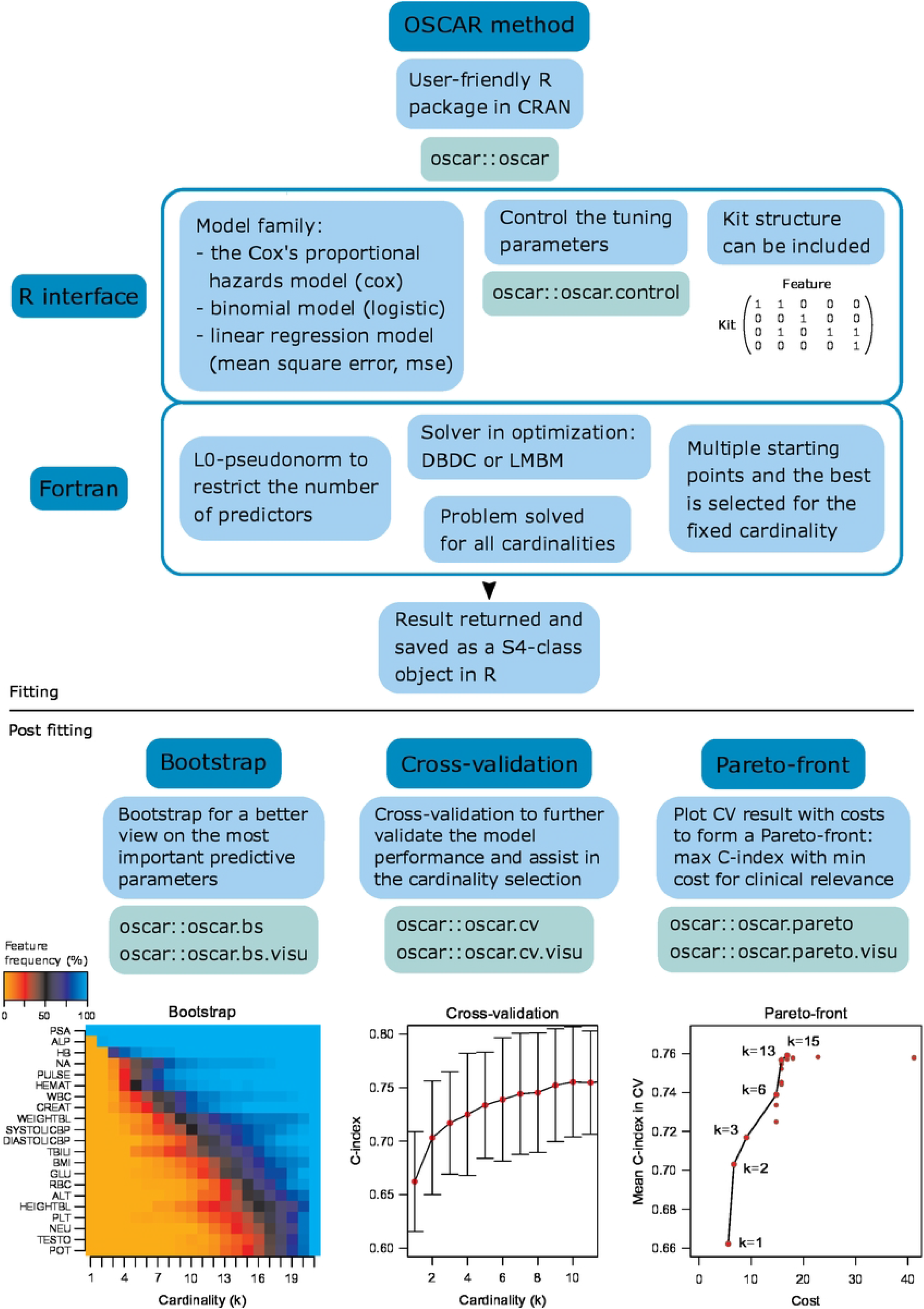
Schematic illustration of the OSCAR method.

## Materials and methods

### Model and algorithm

#### Cox’s proportional hazards model

Our modelling interest is mainly in the patient survival prediction, where we investigate the relationship between features (see Data section) and survival time (overall survival or progression free survival). In the general form, this type of data can be stated as a set

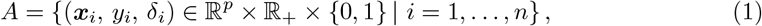

where *n* is the number of observations, ***x***_*i*_ ∈ ℝ^*p*^ is the vector of *p* features, *y_i_* ∈ ℝ_+_ is the observed time and *δ_i_* ∈ {0,1} is the label (value 1 indicates an event and value 0 right-censoring). In addition, we let *t*_1_ < *t*_2_ < … < *t_m_* be increasing list of *m* unique failure times, and *D_i_* be the set of indices of observations failing at time *t_i_* meaning that ties are also allowed to happen.

Survival prediction is traditionally done using Cox’s proportional hazards model [10]. The *hazard* for the patient *i* at time *t* is given with the formula

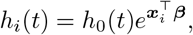

where *h*_0_(*t*) is a shared baseline hazard and *β* ∈ ℝ^*p*^ is an unknown coefficient vector. Our aim is to estimate this vector *β* by maximizing the Breslow approximation of the *partial likelihood* (see [22]). In the following, we denote by 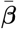 the solution yielding the maximum value for the likelihood.

Instead of maximizing the partial likelihood directly, it is also possible to maximize the scaled log partial likelihood, since this leads to an equivalent solution [5]. This modification gives the *scaled log partial likelihood* of the form

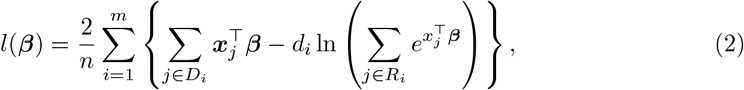

where *R_i_* = {*j*: *y_j_* ≥ *t_i_*} is the set of indices at risk at time *t_i_* and *d_i_* = |*D_i_*| is the number of failures at time *t_i_*. The function –*l* is convex, since it is a sum of linear and log-sum-exp functions [23]. Therefore, instead of maximizing the concave function *l,* it is equivalent to minimize the convex function — *l*. In the following, we concentrate on solving the minimization problem

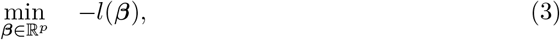

whose solution 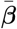 also maximizes (2).

#### Restricting the number of single features

In many real-world applications, the sparsity of the solution for the partial likelihood function is a preferred feature. To favour sparse solutions, a regularization term is typically added to the optimization problem. For example, the elastic net penalization is used in [5], combining *L*_1_- and *L*_2_-norms. In particular, approaches relying on the *L*_1_-norm ensure sparsity to a certain extent.

In our approach, the sparsity of the solution is obtained by using the cardinality constraint to restrict the number of nonzero coefficients in the vector *β*. Thus, the strength of this approach is that it provides us an effective tool to seek solutions with the predetermined model complexity. Instead of considering each feature separately, we may also want to link some features together, if they are always measured together, (i.e. they belong to the same measurement kit). Therefore, we also generalize the cardinality constraint -based subset selection to a case where we restrict the number of selected kits (see Supplementary file S1 File Section 1 for restricting the number of selected kits, instead of single features).

For any vector *β* ∈ ℝ^*p*^, the *L*_0_-pseudonorm ||*β*||_0_ calculates the number of nonzero components. However, it is worth noting that the *L*_0_-pseudonorm is not a proper norm since it is not homogeneous [24], thus the name pseudonorm. In addition, this pseudonorm is discontinuous and nonconvex, making the optimization problem more challenging [15,25].

In the problem (3), sparsity can be achieved by fixing the number of nonzero coefficients *K* ∈ {1,…, *p*} and adding a *cardinality constraint* ||*β*||_0_ ≤ *K*. This results in the following *cardinality-constrained problem*

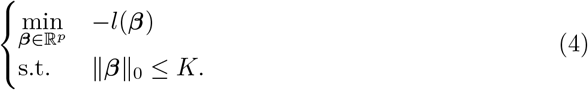

It is known that this problem is difficult to solve due to the combinatorial nature of the constraint, which is also discontinuous. To overcome the discontinuity, we use the approach presented in [15] utilizing the largest-*k* norm to obtain an exact continuous representation of the constraint.

The *largest-k norm* of a vector *β* ∈ ℝ^*p*^ is the sum of the *k* largest absolute value elements:

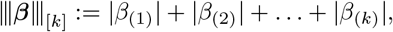

where *β*_(*i*)_ is the element whose absolute value is the *i*-th largest among the *p* elements of *β*. The largest-*k* norm is a proper norm. In addition, it is convex and the constraint ||*β*|_0_ ≤ *K* is equivalent with the constraint ||*β*|_1_ – |||*β*|||_[*K*]_ = 0 [15,25], where ||*β*||_1_: = |*β*_1_| + |*β*_2_| + … + |*β_p_*| = |||*β*|||_[*p*]_ Thus, the problem (4) can be rewritten as

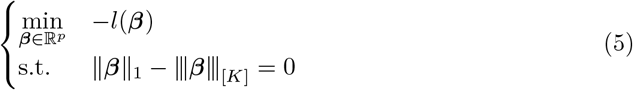

and we have a continuous constraint instead of a discontinuous one. Note that both problems (4) and (5) have exactly the same feasible set. However, the combinatorial structure of the cardinality constraint causes the continuous constraint to be nonconvex. For this reason, the problem (5) may have multiple local solutions and identifying a global or near global solution requires a sophisticated optimizer.

Another disadvantage of the problem (5) is that we still have a constraint. Similarly to [15], we can utilize the penalty function approach [26,27] to rewrite the constrained problem (5) as an unconstrained one

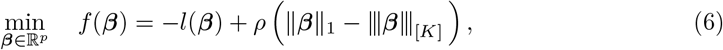

where *ρ* > 0 is a positive penalization parameter. In this reformulation, we are balancing between feasibility and optimality. By selecting a too small value for the parameter *ρ* we do not obtain a feasible solution for the original problem (5). However, by selecting a suitably large value for *ρ*, we have a heavy cost for cardinality constraint violation and end up with a feasible solution. Note that the parameter *ρ* should not be too large since otherwise the penalty term dominates the objective function and we do not obtain an optimal solution for the objective of the constrained problem (5). For this reason, as is typical for penalty function methods, we need to solve the problem (6) sequentially for a series of increasing values of the parameter *ρ* until suitably large parameter value is reached forcing the original constraint in (4) to hold. In practice, this search is done by using a *ρ* value-grid.

One major benefit of the formulation in (6) is that, although its objective *f* is nonconvex and nonsmooth (i.e. nondifferentiable), it is also a *DC function* (Difference of two Convex functions). This means that *f* can be represented in the form *f* = *f*^1^ – *f*^2^ with convex functions *f*^1^ and *f*^2^. This way we can better control the nonconvexity than in the general case. In addition, these convex functions can be selected, for example, as

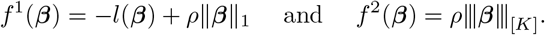

Another interesting aspect of the penalized reformulation (6) is that it can be seen as a modification of the *L*_1_ norm based penalization since the only difference is the largest-*k* norm term – *ρ*|||*β*|||_[*K*]_. Note that this is the term restricting and controlling the upper bound for the number of nonzero features in the problem.

#### Method OSCAR

In this section, we introduce the new algorithm OSCAR (Optimal Subset CArdinality Regression) to solve the cardinality-constrained problem formulated in (4). Since the considered problem is nonconvex, it is well-known that the determination of a global solution is a challenging task, since we may have many local optima and we lack easily verified conditions guaranteeing the global optimality. Due to this, the goal of our new local optimization framework is to find good enough solutions which are close to the global optima. To achieve this goal, our method combines first time the penalty function approach and the double bundle method (DBDC) [16,17] for DC optimization together with an incremental type of an approach to solve the original problem.

OSCAR methodology is designed so, that it does not depend on the specific optimization method, if they are capable of handling both nonsmoothness and nonconvexity. Therefore, our method generalizes beyond DBDC, although it is offered as the default choice. Due to this, we have also incorporated to the R-package of OSCAR the possibility to use the limited memory bundle method (LMBM) [18,19]. LMBM is designed for general nonconvex nonsmooth optimization problems, with the drawback, that it does not benefit from the DC structure of the objective. The most important feature of LMBM is that it scales towards large-scale problems.

As presented above, the first step in OSCAR is to use the penalty function approach to change the rewritten constrained problem (5) to an unconstrained one. Since the objective of the unconstrained problem (6) is DC, we can utilize the DBDC method for the DC optimization to solve it. This enables us to take advantage of the DC structure, since the selected bundle method constructs a nonconvex DC cutting plane model (i.e. an approximation of the objective function, which incorporates both the convex and the concave behaviour of the problem). Another option to solve the problem (6) is LMBM described above.

However, since DBDC and LMBM are only local methods, the quality of solutions for a nonconvex problem strongly depends on the choice of starting points. For this reason, the algorithm OSCAR combines the DBDC and LMBM methods with an incremental type of an approach to generate starting points with higher likelihood of leading to promising parts of the search space. The idea in our incremental approach is to start with solving the cardinality-constrained problem, where only a single predictor (or kit) is allowed to be used initially, and then to increase the number of predictors (or kits) one at a time until the maximal number of predictors is achieved. In particular, we utilize the solution of the cardinality-constrained problem with *i* — 1 predictors to derive promising starting points to the next cardinality-constrained problem with *i* predictors. Since this type of process may end up in a local optimum, we alleviate this challenge via the use of multiple starting points to obtain solution candidates for the problem with *i* predictors.

The OSCAR method is presented in Algorithm 1 for the case where each predictor is considered separately. See Supplementary file S1 File Section 1 for modifications needed with a kit structure. As an input, one needs to give the number of predictors *K*, defining how many predictors maximally can be chosen in the densest cardinality-constrained problem. As an output, the method provides incrementally a solution to each cardinality-constrained problem with *i* predictors for *i* = 1, …, *K* and, thus, we obtain as a by-product a solution also for each cardinality-constrained problem with a smaller number of used predictors (or kits). This means that one can control how many different sparse solutions are generated. Naturally, it is also possible to select *K* = *p*, in which case the problem (5) is solved for all possible numbers of predictors.

**Algorithm 1:**
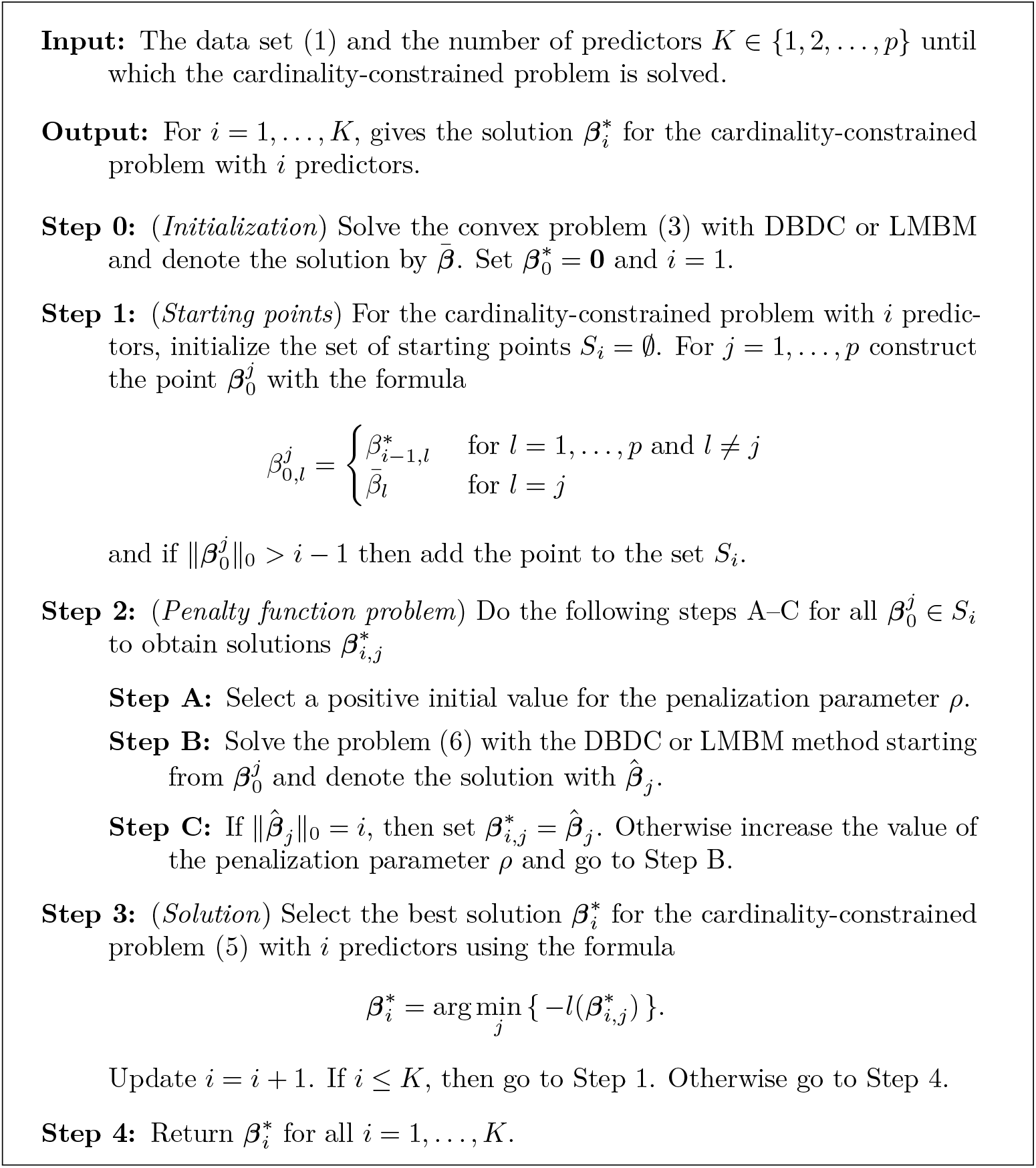
OSCAR.

In Step 1 of Algorithm 1, starting points are generated by varying the previous solution 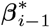 with the best solution 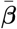 of the scaled log partial likelihood obtained without any regularization. Specifically, in a starting point 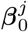 the base is 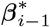, and then we simply substitute predictor *j* with the corresponding value in 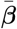. In this way we can easily vary the previous solution but still maintain its main predictors. Note also that each starting point having *i* — 1 predictors is omitted and we only keep the starting points with *i* predictors.

In Step 2B of Algorithm 1, we always use the original starting point. The reason for this is that if the parameter *ρ* is too small, then we may end up with a solution where nearly all the coefficients are nonzero, and therefore, lose the information provided by the original starting point. To avoid such solutions, we do not change the starting point, but instead update the parameter *ρ* until we obtain a solution with the acceptable number of predictors (or kits). This guarantees that the obtained solution does not diverge too much from the previous solution and maintains its best predictors. In addition, this way the method does not become too sensitive to the selection of *ρ*, since too small values of *ρ* are basically omitted.

### Data

For testing the new algorithm for survival prediction, we used one prostate cancer cohort from real-world hospital registry data and three prostate cancer cohorts from randomized clinical trials (see Supplementary Table S1 Table). The patient features were also considered by the clinical examination groups (kits), in which they are measured in clinical practice. Prices for the examinations were provided by the Helsinki University Hospital. The real prices were converted to costs relative to PSA, which was given a reference value of 100. One feature (blood urea nitrogen) without a known cost was ignored. The features are shown in Table 1, along with abbreviations as well as the kit structures and prices.

**Table 1.** Data features.

| Abbreviation | Meaning | Unit | Kit | Price |
| --- | --- | --- | --- | --- |
| AGEGRP | Age group (three groups) | - | Routine measurements | 0 |
| BMI | Body mass index | kg/m<sup>2</sup> |  |  |
| DIASTOLICBP | Diastolic blood pressure | mmHg |  |  |
| HEIGHTBL | Height | cm |  |  |
| PULSE | Pulse | bpm |  |  |
| SYSTOLICBP | Systolic blood pressure | mmHg |  |  |
| WEIGHTBL | Weight | kg |  |  |
| HB | Hemoglobin | g/dl | B-PVKT | 40 |
| HEMAT | Hematocrit | % |  |  |
| PLT | Platelets | E9/l |  |  |
| RBC | Red blood cells | E12/l |  |  |
| WBC | White blood cells | E9/l log |  |  |
| NEU | Neutrophils | E9/l log | TKD | 50 |
| POT | Potassium | mmol/l |  |  |
| ALP | Alkaline phosphatase | U/l log | P-AFOS | 20 |
| ALT | Alanine aminotransferase | U/l log | P-Alat | 20 |
| AST | Aspartate aminotransferase | U/l log | P-AsaT | 20 |
| CA | Calcium | mmol/l | P-Ca | 20 |
| CREAT | Creatinine | umol/l log | P-Krea | 20 |
| LDH | Lactate dehydrogenase | U/l log | P-LD | 20 |
| PSA | Prostate-specific antigen | ng/ml log | P-PSA | 100 |
| TBILI | Bilirubin | umol/l log | P-Bil | 20 |
| TESTO | Testosterone | nmol/l log | S-Testo | 330 |
| NA | Sodium | mmol/l | cB-Het-Ion | 100 |
| MG | Magnesium | mmol/l log | P-Mg | 20 |
| PHOS | Phosphorus | mmol/l log | P-Pi | 20 |
| ALB | Albumin | g/l | P-ALB | 20 |
| TPRO | Total protein | g/l | S-Prot | 20 |
| LYM | Lymphocytes | E9/l log | B-Lymf | 90 |
| CCRC | Calculated creatinine clearance | ml/min log | Pt-GFReEPI | 20 |
| GLU | Glucose | mmol/l log | Gluk | 20 |

Abbreviations, explanations and units for features used in the analyses, as well as kit structures and corresponding prices. Prices were standardized so that PSA has a price of 100.

#### Real-world hospital registry data

The real-world hospital registry data were collected from the advanced prostate cancer patients treated at the Turku University Hospital (TYKS). Patients with castration resistance were selected and data processed as in [9]. Furthermore, only patients with diagnosis of castration resistance dated in 2010 or later were selected, due to the higher sparsity of data in the previous years. In addition, patients with zero or negative survival time or no measurement data were discarded. 195 patients were set aside to be used as an external validation data to evaluate the generalization capability of the model, in order to avoid and assess the risk of over-fitting to the training data. We further eliminated features with over 50% of missing values. All missing data were imputed using median values calculated in the remaining training data (N=590). Median imputation has been previously tested and found adequate [8, 9]. One outlier measurement of systolic blood pressure (>12000 mmHg) was changed into missing before imputation. Patient characteristics for the training data are presented in Supplementary Table S1 Table and the survival curves in the TYKS cohort with respect to the Gleason scores are shown in Fig 2a. The survival curves were as expected, with lowest survival on the highest Gleason scores and highest survival on the lowest Gleason scores. Since cross-variable correlations affect the feature selection process, we investigated these across the available features and present them in Supplementary Fig S1 Fig.

**Fig 2.**
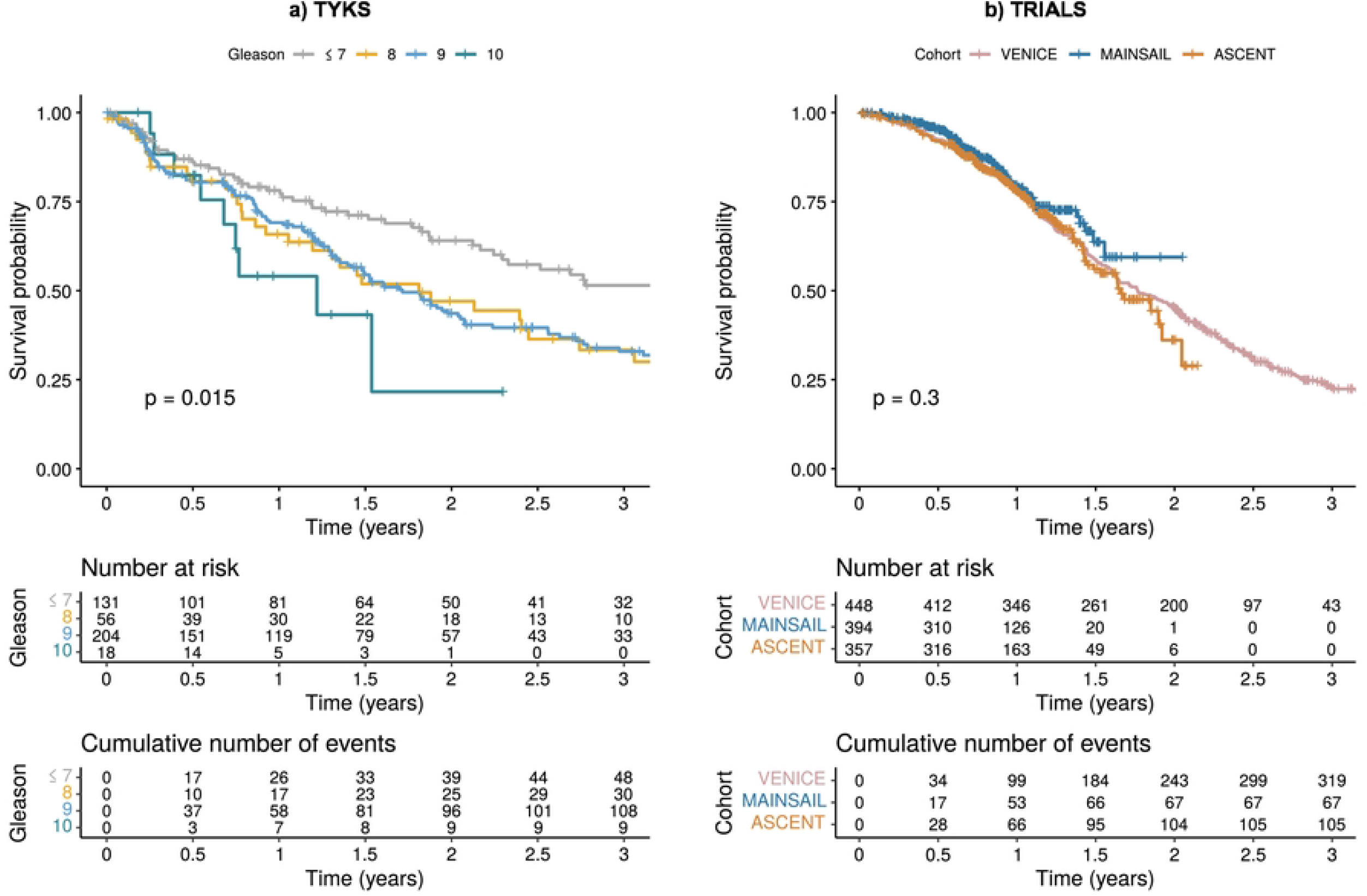
Survival curves. a) Kaplan-Meier survival probability for TYKS patients based on the Gleason scores. b) Kaplan-Meier survival probability for the three trial cohorts: VENICE, MAINSAIL and ASCENT.

#### Randomized clinical trial data

The randomized clinical trial data included in the analyses were previously constructed in the DREAM 9.5 competition (the Prostate Cancer Challenge, PCC-DREAM), hosted by Project Data Sphere (https://www.projectdatasphere.org/). Three prostate cancer patient cohorts, MAINSAIL, VENICE and ASCENT, are included [28–30]. From each cohort, a random set of patients was separated as a validation data (N=132, N=150 and N=119 for MAINSAIL, VENICE and ASCENT, respectively). Features with over 50% of missing values were eliminated. Missing values in each cohort were imputed separately using median values calculated from the corresponding training data sets (N=394, N=448 and N=357 for MAINSAIL, VENICE and ASCENT, respectively). Patient characteristics are presented in Supplementary Table S1 Table and the survival curves per cohort are shown in Fig 2b. The survival curves start similarly, however, the MAINSAIL and ASCENT cohort have a shorter follow-up time. The overall survival trend was also similar to the TYKS cohort (Fig 2a). We also present the correlations between features in the clinical cohorts in Supplementary Fig S1 Fig.

## Results

We investigated the modelling performance of our OSCAR method in four prostate cancer data sets, which portrait two very distinct archetypes of biomedical data. First, we applied the method to advanced prostate cancer cohort obtained from Turku university hospital (TYKS), representing a highly heterogeneous real-world hospital registry cohort. Second, we applied the method to three prostate cancer cohorts obtained from randomized clinical trials, which had been part of a DREAM modelling challenge and had been homogenized previously by the challenge organizers.

The predictive performance was evaluated with concordance index (C-index) [31]. C-index is commonly used in survival analysis as it compares the order of predicted risks to the order of observed survival times [32–34]. To benchmark OSCAR performance, we compared the results to a widely used method LASSO [4], which utilizes *L*_1_-regularization. We also included another *L*_0_-pseudonorm based method APM-*L*_0_ [14], which was chosen based on literature search for *L*_0_-related methods capable of performing survival analysis. We performed CV to assess generalization ability, supported by bootstrapping of the data and subsequent re-fitting of the models to assess robustness of the selected features.

In addition to model accuracy, we evaluated cost-efficiency of the models proposed by OSCAR as a function of feature measurement costs obtained from actual clinical laboratory measurement kit reference costs. We evaluated the model performance of the three methods OSCAR, LASSO and APM-*L*_0_ with respect to the costs calculated with the corresponding number of predictors. This gave us an approximation of the Pareto-front aiming at a good compromise between minimal real-life cost and maximal accuracy, since the underlying problem can be seen as a multi-objective optimization problem of these two objectives.

We also investigated which features were selected as robust predictors. More specifically, we performed BS, in which the model was fitted 100 times to calculate how often (%) each feature was selected as a predictor when a certain cardinality was set. This enabled us to interpret which features are robust predictors that are not sensitive to slight perturbations in the provided data.

### Prognostic prediction for advanced prostate cancer in real-world hospital registry data

Based on the BS evaluation of the OSCAR method in the TYKS cohort (Fig 3a), PSA was the clearly the most robust predictor for overall survival in prostate cancer. However, as can be seen from Fig 3b and c, the original model and CV C-index improved substantially when at least four predictors were chosen. Based on the BS results, the most promising predictors within the explored cardinality values were PSA, hemoglobin (HB), alkaline phosphatase (ALP) and age group (AGEGRP). Notably, the cost remained low when these four predictors were chosen (Fig 3b blue). Adding more predictors did not dramatically improve the OSCAR method accuracy in the training data. However, when more predictors were introduced, creatinine (CREAT) and pulse (PULSE) were chosen for prognostic modelling by OSCAR.

**Fig 3.**
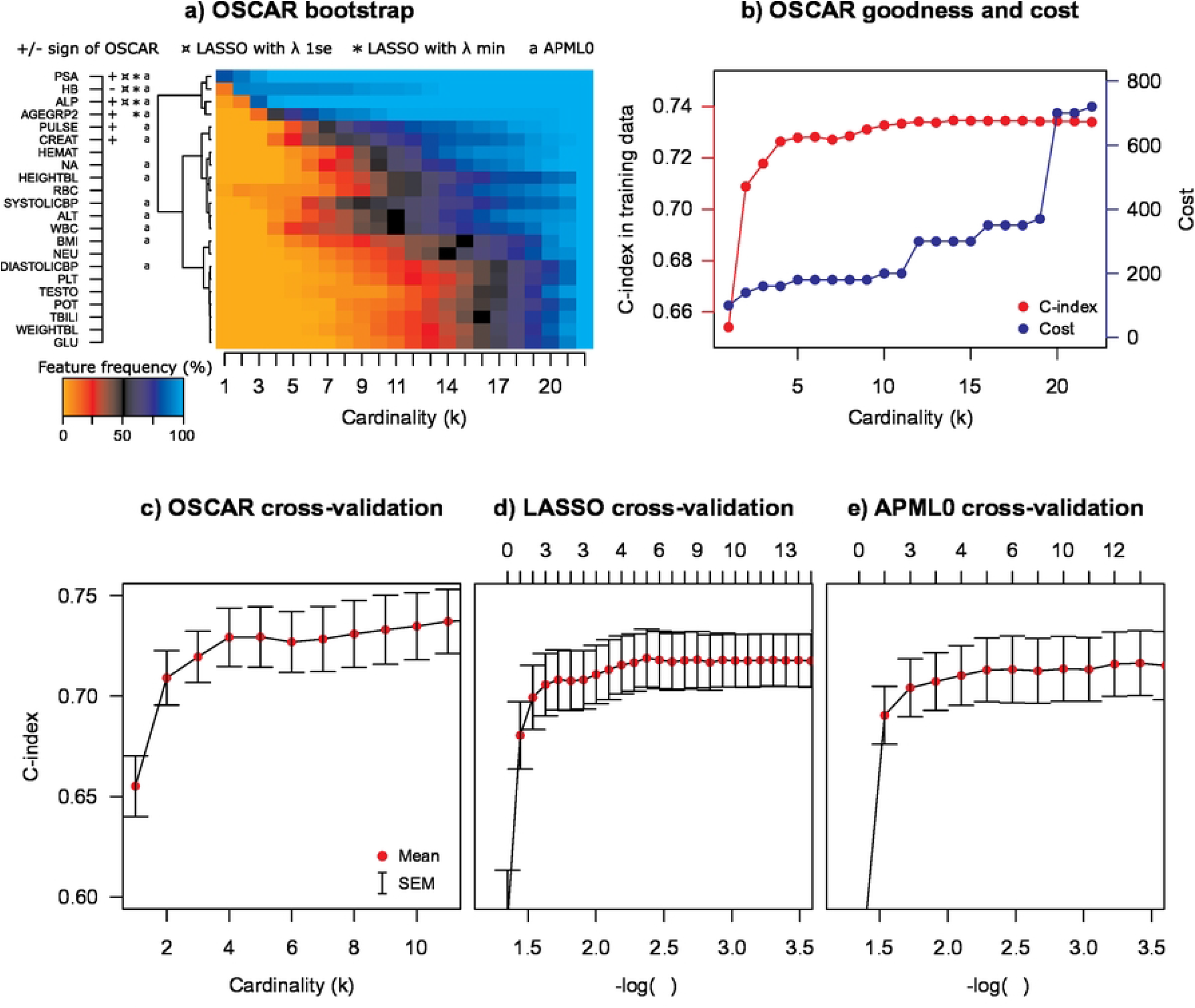
TYKS data: a) OSCAR BS performance. +/− denotes the sign of the coefficient in the model. Positive coefficient: higher predictor value leads towards high risk. Negative coefficient: higher predictor value leads towards low risk. ¤ denotes features selected by LASSO with *λ*_1*se*_ and * denotes features selected by LASSO with *λ_min_*. a denotes features selected by APM-*L*_0_. Color denotes how often among 100 bootstrap runs a feature is selected when a certain cardinality is set (1 meaning 100%). b) OSCAR accuracy in the TYKS training data (C-index), and cost with respect to the allowed number of predictors. Cost is calculated by kits and a kit price is added if any feature from a kit is used. c) CV performance of OSCAR. d) CV performance of LASSO. The numbers at top indicate the number of predictors selected by a specific lambda. e) CV performance of APM-*L*_0_. The red dots denote the mean values and error bars denote the standard errors of mean (SEM) calculated over the CV folds.

In general, OSCAR resulted in improved performance in terms of C-index in CV, when benchmarked against LASSO and APM-*L*_0_ methods (Fig 3c-e). All methods exhibited roughly similar amount of variation over the folds in the CV. Of note, since OSCAR estimates do not shrink toward zero and are instead either included or excluded, which may partly explain the saturation effect in the CV performance curves. Alternatively, in our modelling task the number of predictors (p=22) was relatively low in comparison to the number of patients (N=590). All the three methods selected similar predictors (Fig 3a). For example, LASSO with conservative lambda (*λ*_1*se*_) selected three predictors (PSA, HB and ALP), which are the same as the most important predictors of OSCAR based on the BS. The choice of *λ* (penalization coefficient) in LASSO and APM-*L*_0_ is typically chosen either based on a local optimum for CV performance (*λ_min_*) or when a solution is within a standard error’s range of the local optimum (*λ*_1*se*_). In OSCAR, to avoid arbitrary choices for the crucial model penalization, we leverage the use of bootstrap-based inference to explore feature robustness in addition to the CV generalization ability.

To compare the methods in terms of implementation costs, we investigated the mean C-index in CV of the three methods OSCAR, LASSO and APM-*L*_0_ with respect to the costs calculated with the corresponding number of predictors or lambdas (Fig 4). Interestingly, the Pareto-front for OSCAR CV performance vs. cost suggested multiple candidate models, which could be then refined using the domain-expert based guidance. The models from these approximated Pareto-fronts were subsequently selected for testing in the left-out validation data (Fig 5) to further assess model generalization ability beyond the already observed training data. The observed C-index in the validation data were similar to that in the training data. All three methods performed well in the validation data, with OSCAR slightly better for most of the costs (or number of predictors).

**Fig 4.**
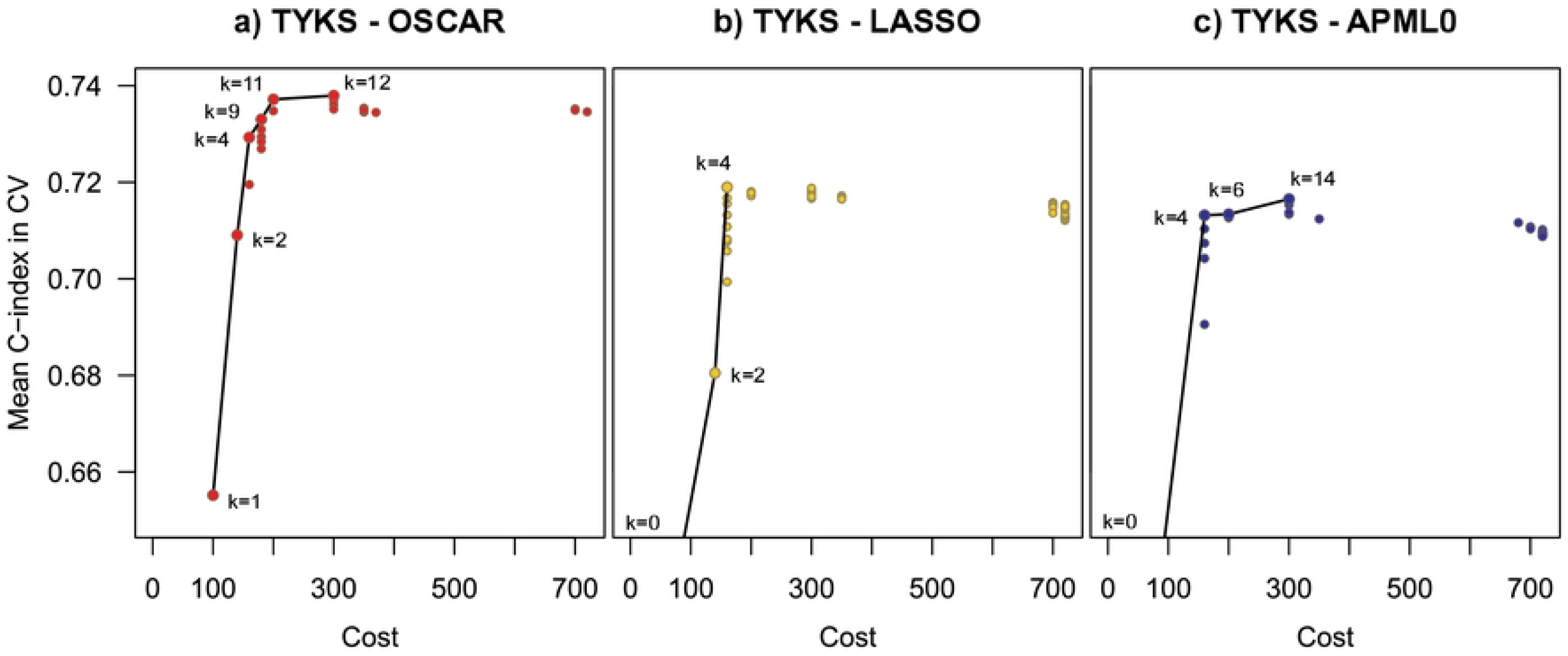
Model accuracy in CV with respect to the cost for a) OSCAR, b) LASSO and c) APM-*L*_0_. The approximated Pareto-front is marked with black line. Number of predictors in each Pareto-point is noted next to the point. The costs were calculated with the corresponding number of predictors (OSCAR) or corresponding lambdas (LASSO and APM-*L*_0_) using predictors chosen in the model fitted for the entire training data (e.g., Fig 3b).

**Fig 5.**
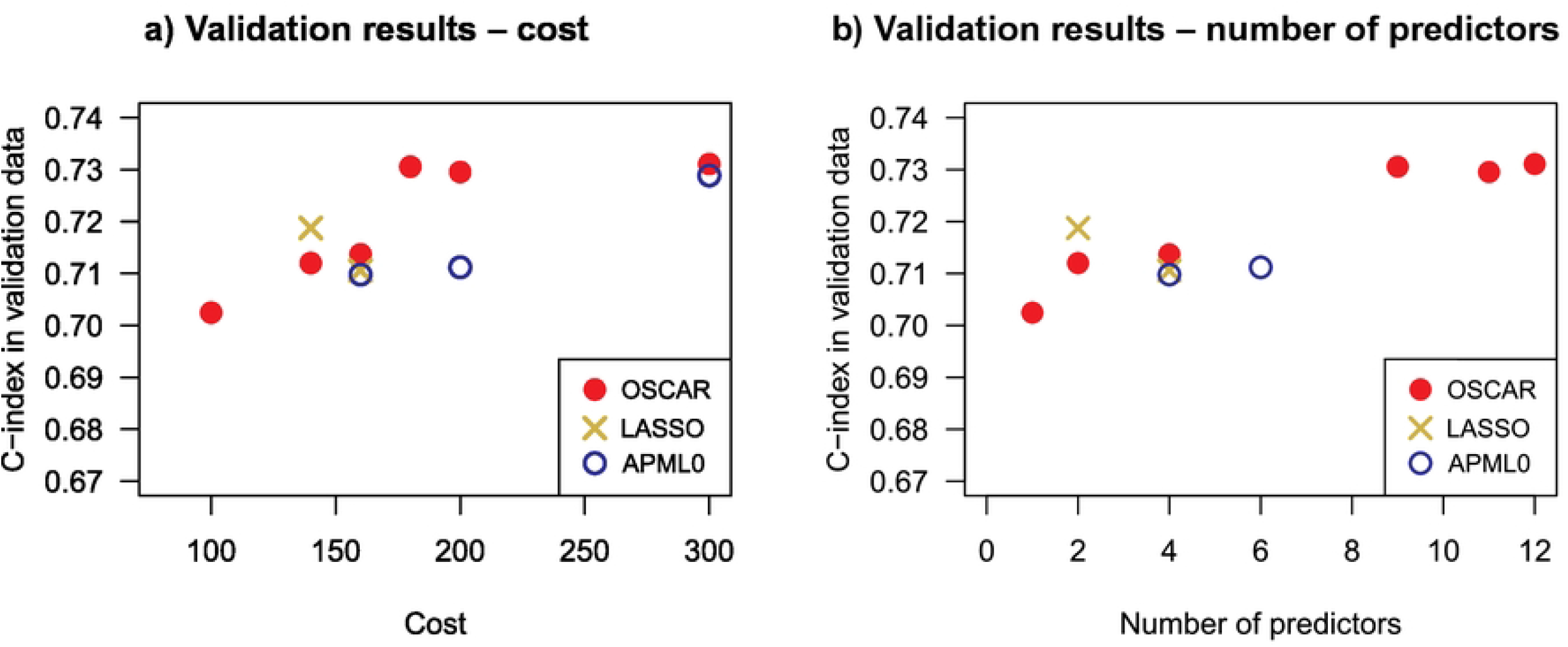
Model accuracy in validation data cohort for OSCAR (red filled circles), **LASSO** (**yellow crosses**) **and APM**-*L*_0_ (**blue hollow circles**) a) with respect to the corresponding costs, b) with respect to the corresponding number of predictors. Only the performance of the models in the corresponding approximated Pareto-fronts are presented.

We further considered scenarios in which the Pareto-front is of no special interest and only a single model prediction is required. For this purpose, the model CV performance was inspected using a smoothing spline fitted on the performance as a function of cardinality. The smoothing spline was investigated to find a shoulder-point (i.e. a point where the model accuracy saturates and no longer improves when more predictors are allowed). In Supplementary Fig S2 Fig, the spline is fitted for the TYKS data models (Fig 3b). The shoulder-point was selected among the points where the second derivative indicated a steep saturation in the curve (i.e. crossing the x-axis). Using this strategy, six predictors (PSA, HB, ALP, AGEGRP, CREAT and PULSE) were selected, suggesting a similar model as previously identified with CV and BS. These results demonstrate that even though the three methods had a trend toward the same features, OSCAR’s generalization ability was similarly good or better than those using shrinkage-based coefficient estimates.

### Prediction with kit structure

While the most typical approach is to choose features one at a time, as presented above, features may be available as groups. In clinical practice, features are often measured together as kits (e.g., complete blood count), and therefore including a single feature from a kit in the model leads to availability of measurements for the rest of the kit’s features as well. As the extra features become available at the same cost, it is economical to consider including all of the kit’s features in the model simultaneously.

Such a kit structure can be easily included in the OSCAR method (see Supplementary file S1 File Section 1), and was investigated in the TYKS data set. Kit structures used in the analysis are presented in Table 1. Consistent with the non-kit version in the previous section, PSA was the most relevant predictor in the TYKS data (Fig 6a). When two kits are allowed, the model suggests B-PVKT (complete blood count), which includes HB, platelets (PLT), white blood cells (WBC), red blood cells (RBC) and hematocrit (HEMAT). While the inclusion of B-PVKT was largely driven by HB, which had been identified as an important predictor in the non-kit approach, four other predictors were now available for model fitting as well. The model fit C-index levels were slightly lower than those with the non-kit prediction. For example, with two kits (total of six predictors), C-index was 0.708 (Fig 6b), whereas the non-kit model of six predictors had C-index on 0.728 (Fig 3b). This is due to trend of including less prognostic features when a kit includes also a highly prognostic feature. However, the cost of six predictors in the non-kit model was 180, whereas the cost for six predictors (two kits) in the kit structure model was 120.

**Fig 6.**
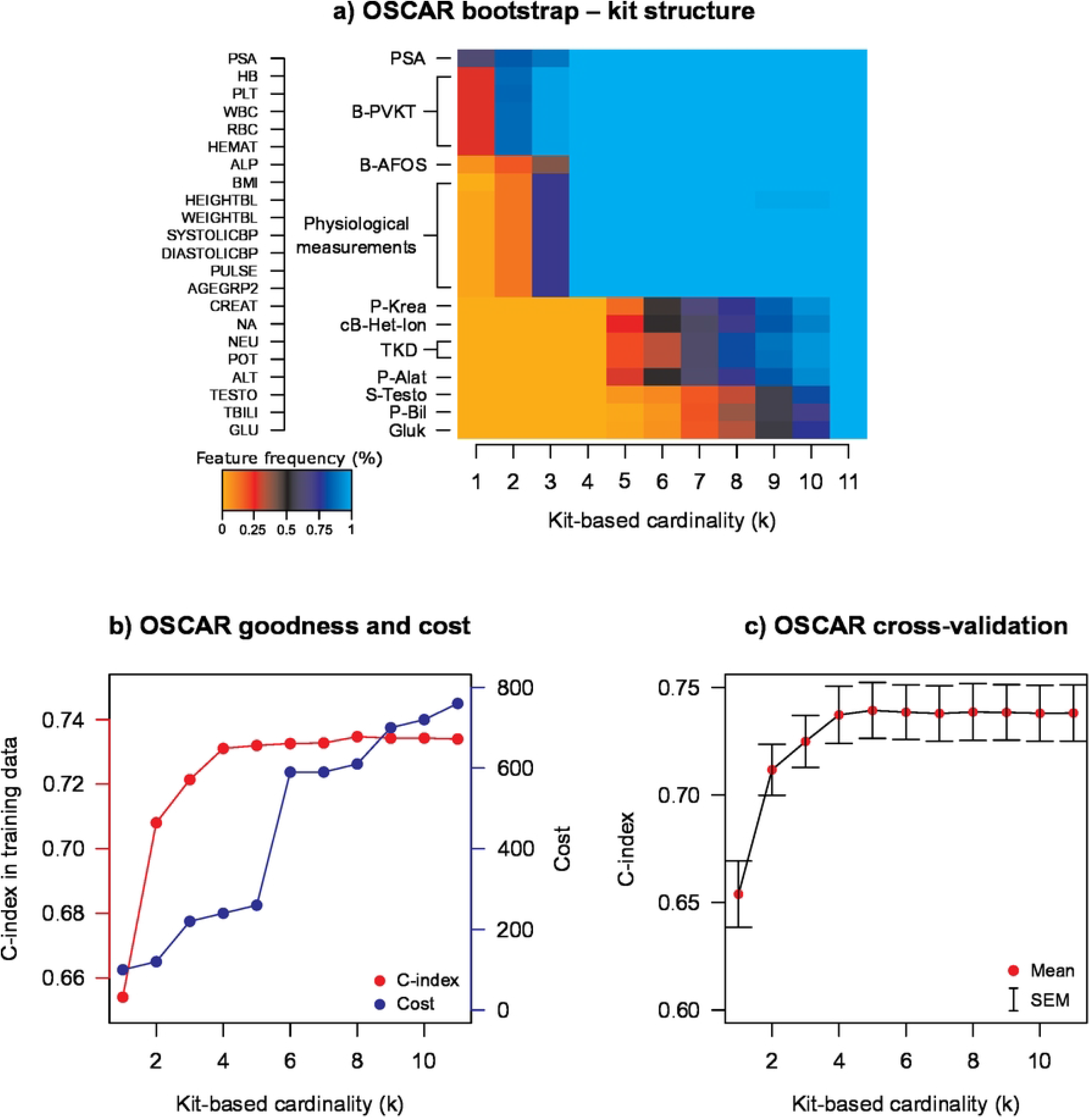
Model performance of OSCAR when kit structure is used. a) Bootstrap performance. b) Goodness (C-index) and cost. c) CV performance. The red dots denote the mean values and error bars denote the standard errors of mean (SEM) calculated over the CV folds.

The overall levels of C-index in the CV were similar with or without the kit structure (Fig 6c), when compared to the non-kit prediction (Fig 3c). With the kit structure, the model included features that would not be likely picked by the non-kit model, such as the above mentioned PLT, WBC and RBC. Furthermore, more parameters could be included while keeping the cost low. For example, with two kits, the cost was 120, when including six parameters, whereas without the kit structure, a higher cost was paid with only two parameters. However, with more parameters, the risk of overfitting increases. These results demonstrate how the OSCAR method enables the inclusion of clinically relevant kit structures and addition of multiple model predictors at a given cardinality. In the presented application, the models retained a similar level of generalization ability regardless whether or not the kit structure was taken into account.

### Prognostic prediction for prostate cancer patients in clinical trial data

To investigate how the developed methodology would perform in a more systematically collected and homogenized clinical cohort, we investigate the model performance in three clinical trial data cohorts. One of the striking differences was, that in contrast to the real world cohort TYKS, PSA was significantly less prognostic factor in the three trial data cohorts. In the ASCENT cohort, PSA was distinguished as a prominent predictor (Fig 7 bottom row); however, if only one predictor was allowed, ALP was selected most often in the BS analysis. Furthermore, ALP was selected as the main predictor in the VENICE cohort (Fig 7 top row). In the MAINSAIL cohort, ALP was not detected as a prognostic feature (Fig 7 middle row), and instead, lactate dehydrogenase (LDH) was the most prominent predictor. In the VENICE and MAINSAIL cohorts, HB was selected most often as the second predictor.

**Fig 7.**
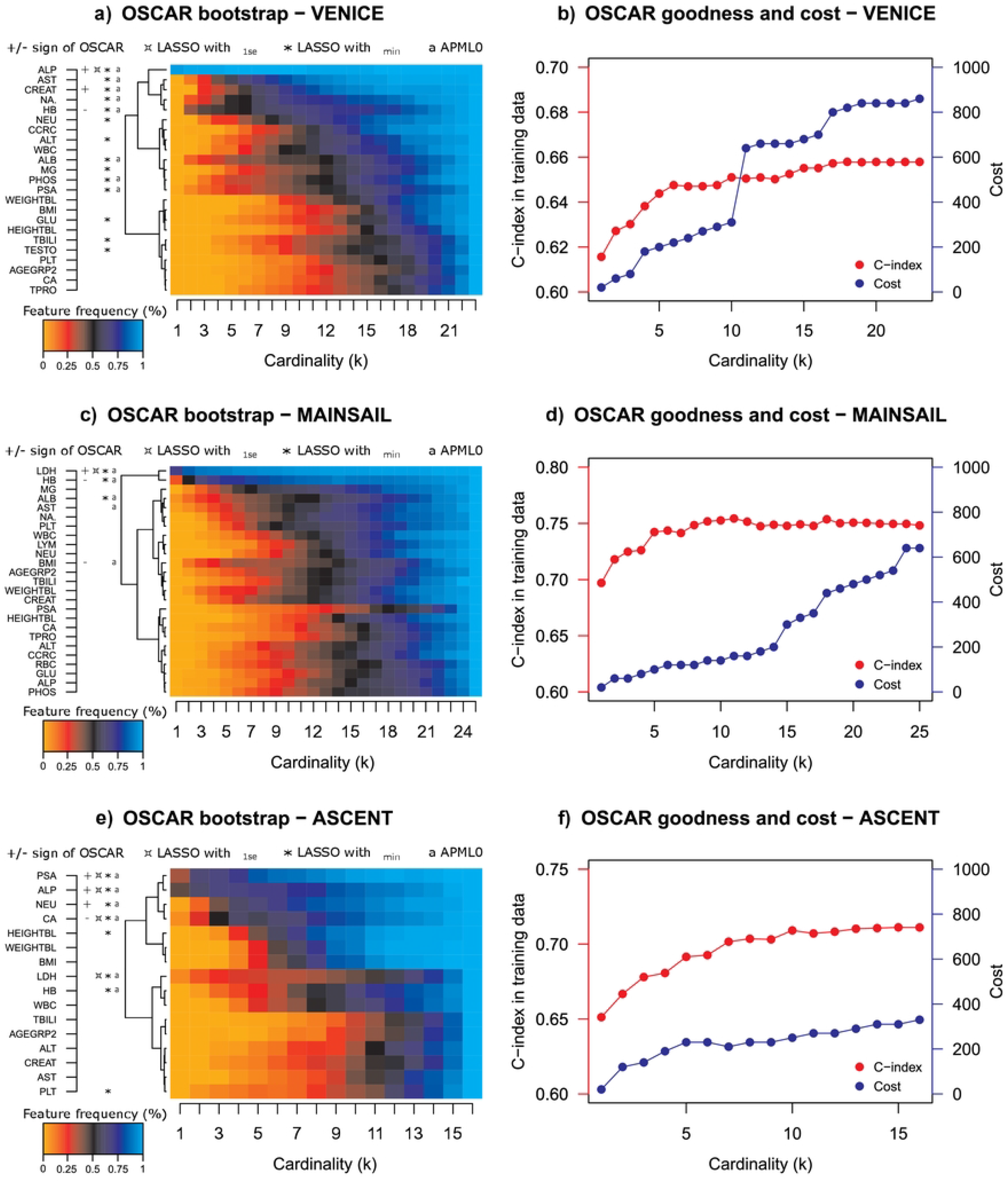
Left panel: BS plots for three trial cohorts. Right panel: Model goodness (C-index) and costs with respect to allowed number of predictors. Cost is calculated by kits and a kit price is added if any feature from a kit is used.

ALP and HB were also highly prognostic in the real-world TYKS cohort. Unfortunately, the otherwise highly interesting LDH was not available in the TYKS cohort due to high percentage of missing values (>80%, Supplementary Table S1 Table). Similarly, TYKS data was missing aspartate aminotransferase (AST), which had notable prognostic power in the VENICE cohort. We note that AST was also, along with LDH, ALP and HB, detected as one of the most important predictors in the original DREAM 9.5 Prostate Cancer Prediction Challenge [6]. The lack of PSA as the clear top-predictor is also in line with the DREAM 9.5 challenge results, rather multiple predictors and their interactions need to be considered for maximal prognostic accuracy. Furthermore, PSA’s elevated prominence as a prognostic predictor may be also biased by data generation and reporting, as it is routine measured in prostate cancer follow-up, while real-world clinical applications may be less prone to adapt novel markers into routine use.

In the VENICE cohort, after selection of these main predictors that appeared over all the trial cohorts, it became less clear which features had most prognostic power on patient survival. However, based on the model accuracy and the CV results (Fig 7 and Fig 8 top rows), a higher model accuracy was reached with additional predictors. Potential candidate features that improved model performance were AST, CREAT, sodium (NA), HB, and albumin (ALB). In the CV analysis, the OSCAR method resulted in higher mean C-index than LASSO and APM-*L*_0_ (Fig 8 top row). However, all the three methods suggested similar predictors, indicating their importance and robustness.

**Fig 8.**
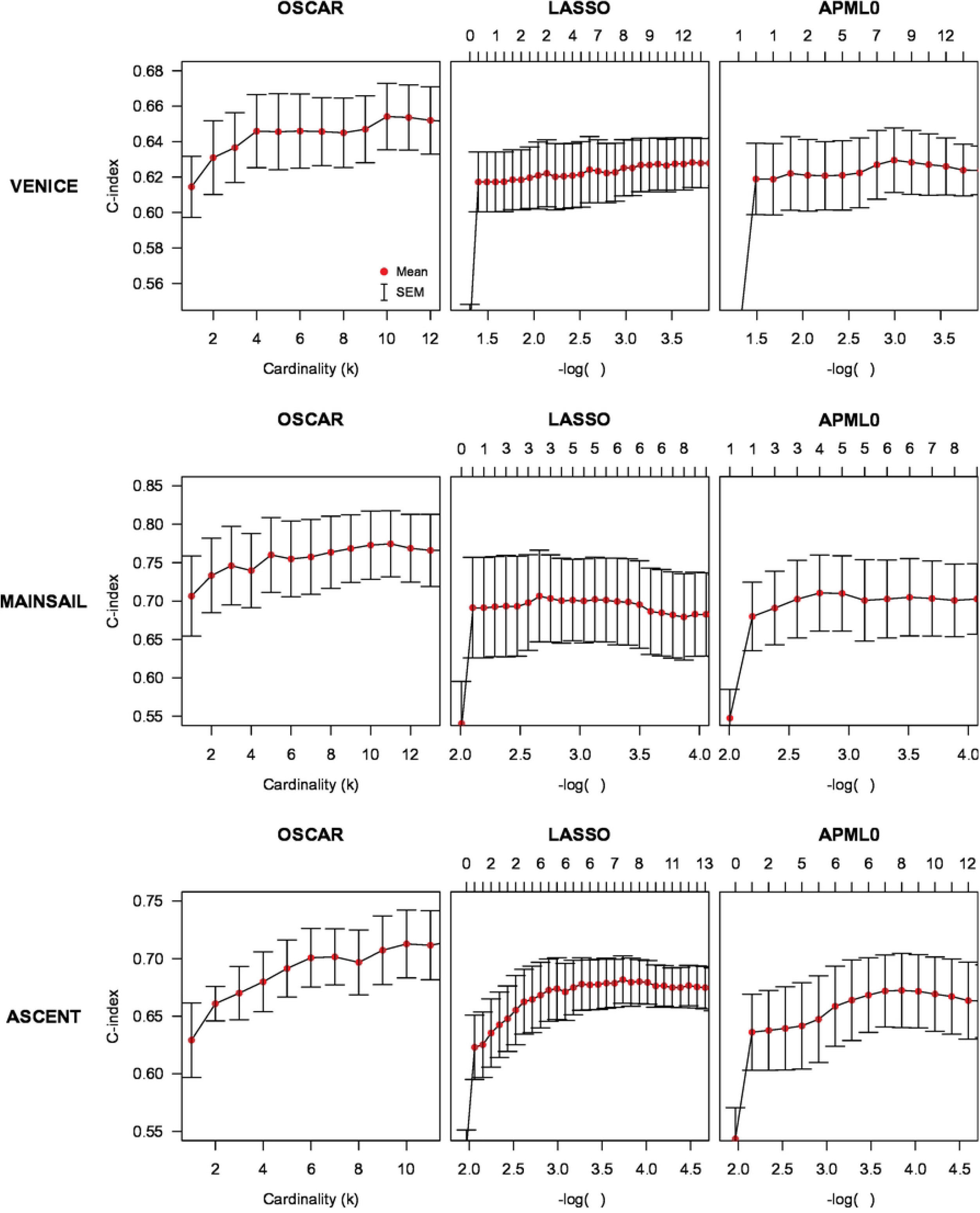
Left panel: CV performance of OSCAR in the three trial cohorts. Middle panel: CV performance of LASSO in the three trial cohorts. Right panel: CV performance of APM-*L*0 in the three trial cohorts. The red dots denote the mean values and error bars denote the standard errors of mean (SEM) calculated over the CV folds.

In the MAINSAIL cohort, a relatively high C-index was reached by roughly five predictors, and adding more predictors did not considerably increase the C-index. In the CV, a local maximum was also reached with three predictors (Fig 8 middle row). Thus, based on the BS analysis, in addition to LDH and HB, features like magnesium (MG), body mass index (BMI), ALB, AST and weight (WEIGHT) were suggested as potential candidates. When compared to LASSO and APM-*L*_0_, OSCAR again resulted in higher mean C-index (Fig 8 middle row).

In the ASCENT cohort, PSA and ALP were the most important predictors (Fig 7 bottom row). Allowing more predictors, such as neutrophils (NEU), calcium (CA), LDH and HB, further increased the C-index. Similarly to the other clinical trial cohorts, OSCAR resulted in the highest mean C-index in the CV analysis when compared to LASSO and APM-*L*_0_ (Fig 8 bottom row).

To investigate the implementation costs, the mean CV accuracies were inspected with respect to the cost in all three trial data cohorts and for all three methods (OSCAR, LASSO and APM-*L*_0_) (Supplementary Fig S6 Fig). For each of the cohort-method pairings, the approximated Pareto-fronts were analyzed. Similarly to TYKS data, OSCAR method resulted in higher accuracies when compared to LASSO and APM-*L*_0_ at the same cost levels. Next, the models corresponding to the approximated Pareto-fronts were applied in the validation data set (Fig 9). In the validation data, the models may have exhibited some overfitting as the highest validation C-index was often reached already with a relatively low feature cost. In general, OSCAR performed well in validation considering the objective of simultaneously maintaining high validation C-index and low cost. Ultimately, a feasible compromise between validation performance and clinical cost would then rely on the domain expert’s decision making.

**Fig 9.**
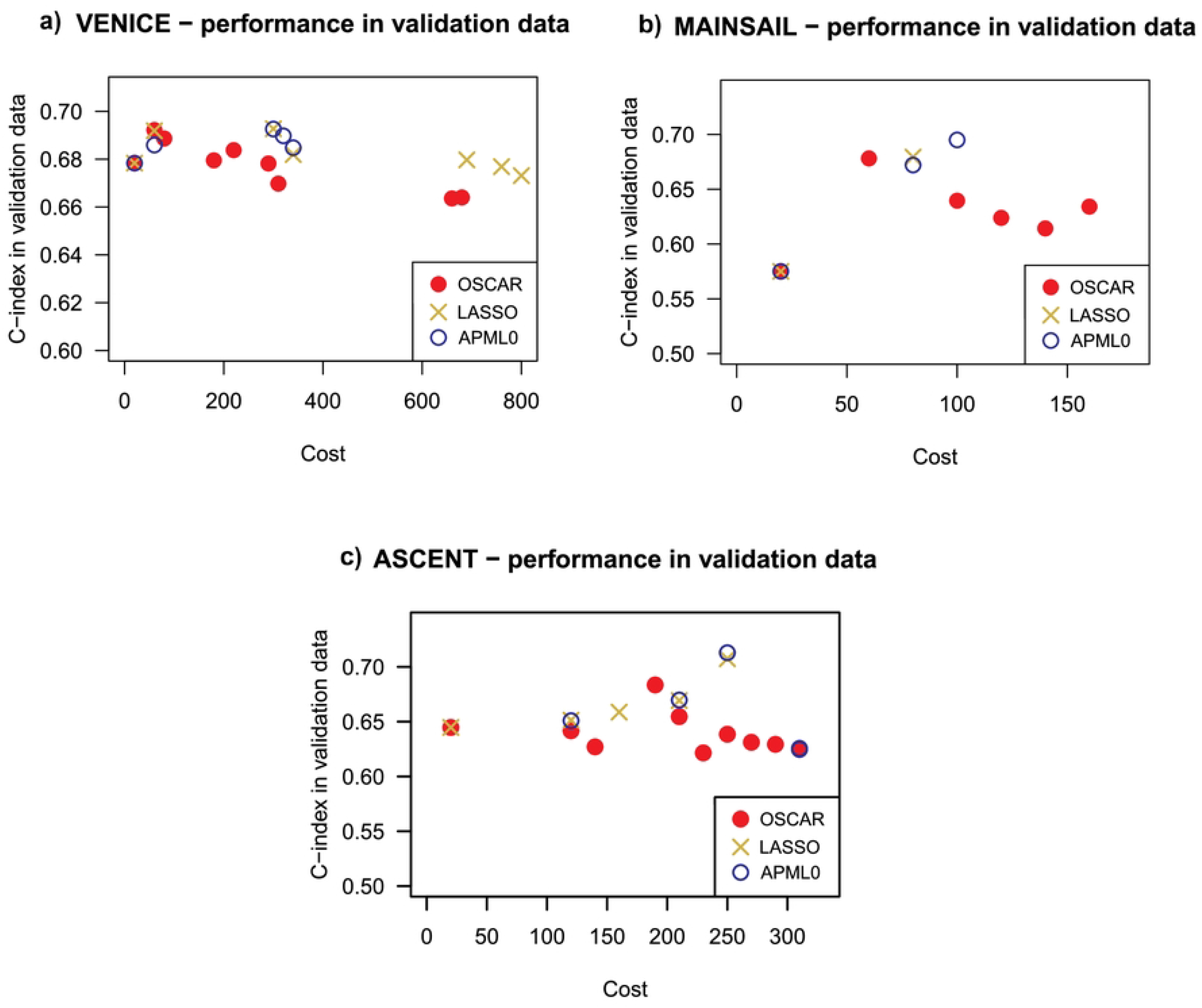
Model accuracy in validation data cohort for OSCAR (red filled circle), LASSO (yellow diamond) and APM-*L*_0_ (blue hollow circle) in the three trial data cohorts VENICE, MAINSAIL and ASCENT. Only the performance of the models in the corresponding approximated Pareto-fronts are presented (see Supplementary Fig S6 Fig).

Based on the spline fittings and its derivatives for VENICE (see Supplementary Fig S3 Fig), OSCAR selected three predictors (ALP, HB and CREAT). For the MAINSAIL cohort (see Supplementary Fig S4 Fig), OSCAR selected three predictors (LDH, BMI and HB). For the ASCENT cohort (see Supplementary Fig S5 Fig), OSCAR selected four predictors (PSA, NEU, ALP and CA).

These results demonstrate that the models based on the trial cohorts slightly differ in terms of the selected model parameters from each other, and also from the real-life cohort TYKS. However, some differences may be caused by the lack of data in some of the cohorts (e.g., LDH lacking from TYKS). The three compared methods selected similar predictors within a cohort. However, the OSCAR method improved the prediction accuracy in training data without increasing the cost.

## Discussion

In this work, we have introduced a new *L*_0_-regularized regression methodology OSCAR, and demonstrated its use in the context of prostate cancer survival prediction both in real-world hospital registry and clinical cohort data. The OSCAR method utilizes *L*_0_-pseudonorm as a penalty term to restrict the number of predictors. Unlike typical approaches trying to tackle *L*_0_ pseudonorm’s difficult formulation, OSCAR restructures the problem so that no approximation is required and the original solution can be then optimized in an exact manner. Since the pseudonorm is discontinuous and nonconvex, the optimization problem becomes NP-hard and computationally heavy [11]. In the OSCAR method, the *L*_0_-pseudonorm based penalty was rewritten for easier management, and this leads to a regularization term in the form a DC (Difference of two Convex functions) composition. The optimization was done using DBDC algorithm [16,17]. This is more sophisticated and more suitable for nonconvex problems than, for example, the classical coordinate descent. DBDC was supplemented by a more computationally efficient optimizer LMBM.

We compared OSCAR to LASSO, a widely used method in survival prediction, and APM-*L*_0_, a *L*_0_-based survival prediction method [14]. All three methods selected similar predictors. In general, OSCAR was the optimal choice based on the CV analyses. This is partly because the *L*_0_-pseudonorm allows the model coefficients to vary freely from zero, unlike in, for example, LASSO, which pushes the coefficients towards zero. LASSO and APM-*L*_0_ utilize the coordinate descent in optimization, which are more prone to local optima when compared to the DBDC algorithm. Despite the *L*_0_ approach, APM-*L*_0_ performed similarly to LASSO, most likely because it incorporates *L*_1_ and *L*_2_.

We investigated the model performance in three data cohorts, one from hospital registry data (TYKS) and three from clinical trials (VENICE, MAINSAIL, ASCENT). In the TYKS cohort, the OSCAR method suggests PSA, HB, ALP and age group as the main predictors. Similar trend is also observed if kit structure was included. PSA reflects the disease severity, especially at disseminated state and in treatment-resistant disease [35]. Thus, PSA has been numerously acknowledged as an important predictor for prostate cancer, and it is used in practice to determine and monitor the state or occurrence of prostate cancer. PSA’s elevated prominence as a prognostic predictor in our hospital registry data may thus be biased by data generation and reporting. High level of ALP is associated with metastases in advanced prostate cancer and it is also measured in the clinical practice to monitor the spreading of cancer into the bones [36]. Metastases typically lead to decreased survival time and, thus, an increased risk of death, therefore predictors associated with metastases have an intuitive explanation as to why they have prognostic power. HB is generally a good indicator of a person’s health. Similarly to HB, age group is linked to the overall health of a person as overall disease burden is typically higher and physical performance status is lower. Since we predict overall survival, higher age leads to decreased survival time regardless of the cancer related characteristics, which somewhat complicates its survival interpretation.

In the VENICE cohort, ALP prevailed as the most prominent predictor, and AST, CREAT, NA, HB, and ALB followed as additional predictor candidates. As mentioned above, ALP is associated with metastases and thus poor prognosis. AST tests for liver damage, and it has been associated with multiple cancers including prostate, bladder, testicular and small cell lung cancer [37–41]. CREAT is related to kidney malfunction, and NA metabolism also mainly reflects kidney function. Taken together, these prognostic factors therefore reflect potential organ failure or organ damage burden. As such, their use in prognostic models are highly justified and intuitive.

Albumin is a protein that maintains fluid balance and osmolality in bloodstream and it is associated with malnutrition and problems in intake of nutrients in the gut [42]. Compromised intake of nutrients may be caused by cancer, cancer-related decrease in daily performance or cancer treatments, suggesting a potential link between ALB and cancer prognosis [43, 44]. In addition, ALB is considered to reflect liver function and in metastasized, castration-resistant prostate cancer, and lowered levels of ALB is known to associate with increased tumor burden [45, 46].

In the MAINSAIL cohort, HB and ALB were again identified as notable prognostic features. In addition, LDH was selected systematically in the BS analysis as a key predictor. LDH is an enzyme participating in energy production in nearly all tissues. Damaged tissues release LDH, which has been linked to cancer burden [43].

Similarly to the VENICE cohort, AST was among the top predictors in the MAISAIL cohort. In addition, BMI, MG, and WEIGHT had considerable prognostic power, of which MG is a pivotal part of metabolism.

In the ASCENT cohort, similar features were selected consistently in the BS analysis: PSA and ALP, along with NEU, CA, LDH and HB. NEU are white blood cells that kill bacteria and help in wound healing. They have also been associated with cancer, despite the previous belief of neutrality against cancer [47, 48]. Especially advanced cancer accumulates NEU, which therefore becomes a predictor of poor survival. Unlike in other two trial cohorts, CA was selected among six top predictors in the ASCENT cohort. CA is a mineral especially involved in bone metabolism. Since the prostate cancer is prone to metastasize in bones, the CA balance may be affected by the cancer development. However, another causation could also be considered since high calcium intake has been associated with increased risk of advanced prostate cancer [49,50].

Taken together, there are still some potential improvements to be considered, despite the already promising validation results with comparable accuracy and reasonable model parameters. Due to the inclusion of *L*_0_-pseudonorm, the optimization problem becomes NP-hard and computationally heavy. Thus, further development of the optimization process, such as using different optimization algorithms or refining the selection of starting points, could potentially improve the running time and model solution. For example, the coordinate descent is a naive but extremely computationally lean optimizer, and it could be considered as a potential alternative in the future work complemented by suitable heuristics. Another development possibility is to reformulate the objective function to take into account also the user-provided costs of features and kits. However, this will lead to a discrete optimization problem.

## Conclusion

We have explored and made available a novel approach to the *L*_0_-regularized regression, which has previously gone under-represented within the domain of regularized regression partially due to challenges related to solving the discrete optimization task. Our approach is exact to *L*_0_-penalty as it does not utilize any approximation of the *L*_0_-pseudonorm, but instead uses its exact DC (Difference of two Convex functions) reformulation, bringing the optimization task to the continuous domain. In addition, we have incorporated the kit structure into the method, enabling the selection of features as groups that they are measured in the practice. Since the measurements have a potentially high costs, the model sparsity allows the selection of the most prognostic features to avoid excessive costs by addition of redundant predictors. The costs were investigated along with model accuracy. This gave us an approximation of the Pareto-front based on the minimal cost and maximal accuracy, since the underlying problem can be seen as a multi-objective optimization problem with two objectives: accuracy and cost. The multi-objective optimization could be regarded as a new way of providing models that are highly relevant to real-world applications, rather than merely optimal according statistical metrics. This way the regularized methodology can also leverage domain-expert knowledge in choosing the final suitable model.

The OSCAR method demonstrated efficient performance in the context of metastatic castration resistant prostate cancer in real-world hospital registry data, as well as in the three clinical trial data cohorts. Our results brought insights into best markers, which to some extent differ between real-world registry data and clinical trial data, possibly due to differences in cohort patient characteristics, missingness, or data reporting practices. We benchmarked our methodology against highly popular regularization methods, readily available for R users, such as LASSO, and demonstrated comparable performance of our *L*_0_-approach. The methodology has been implemented and distributed as a user-friendly R-package accompanied by a wide range of useful helper functions and a set of efficient Fortran optimizers called from within the R-package. The OSCAR method is easily accessible through the Central R Archive Network (CRAN).

## Availability

The latest open source git version control consisting of R, Fortran, and C code for the *oscar* package is available at: https://github.com/Syksy/oscar *oscar* R-package is available at the Central R Archive Network (CRAN) at: https://CRAN.R-project.org/package=oscar

Representative simulated real-world registry data are provided within the *oscar* R-package. Access to the TYKS hospital registry data may be requested via the Auria Clinical Informatics unit at the Turku University Hospital. The DREAM 9.5 mCRPC processed clinical cohort data are available from: https://www.synapse.org/#!Synapse:syn4756967

## Supporting information

**S1 File. Supplementary material.**

**S1 Table. Data characteristics for the data cohorts: TYKS, MAINSAIL, VENICE and ASCENT.** Included in S1 File.

**S1 Fig. Correlations between features in each data cohort.** Included in S1 File.

**S2 Fig. Smoothing spline for TYKS cohort.** Included in S1 File.

**S3 Fig. Smoothing spline for VENICE cohort.** Included in S1 File.

**S4 Fig. Smoothing spline for MAINSAIL cohort.** Included in S1 File.

**S5 Fig. Smoothing spline for ASCENT cohort.** Included in S1 File.

**S6 Fig. Approximated Pareto-fronts in trial data cohorts.** Included in S1 File.

## Authors’ contributions

**Conceptualization:** Anni S. Halkola, Kaisa Joki, Tuomas Mirtti, Marko M. Mäkelä, Tero Aittokallio, Teemu D. Laajala.

**Data curation:** Anni S. Halkola, Teemu D. Laajala.

**Formal analysis:** Anni S. Halkola, Kaisa Joki, Teemu D. Laajala.

**Funding Acquisition:** Tero Aittokallio, Teemu D. Laajala.

**Methodology:** Anni S. Halkola, Kaisa Joki, Marko M. Mäkelä, Tero Aittokallio, Teemu D. Laajala.

**Resources:** Tero Aittokallio.

**Software:** Anni S. Halkola, Kaisa Joki, Teemu D. Laajala.

**Supervision:** Teemu D. Laajala.

**Validation:** Anni S. Halkola, Tuomas Mirtti.

**Visualization:** Anni S. Halkola.

**Writing – Original Draft Preparation:** Anni S. Halkola, Kaisa Joki, Tuomas Mirtti, Teemu D. Laajala.

**Writing – Review & Editing**: Anni S. Halkola, Kaisa Joki, Tuomas Mirtti, Marko M. Mäkelä, Tero Aittokallio, Teemu D. Laajala.

## Acknowledgements

The authors would like to thank Mika Murtojärvi for his advice regarding the hospital registry data processing, and Arho Virkki for administrating the TYKS hospital registry data access.

